# Electromyography Classification during Reach-to-Grasp Motion using Manifold Learning

**DOI:** 10.1101/2020.07.16.207639

**Authors:** Elnaz Lashgari, Uri Maoz

## Abstract

Electromyography (EMG) is a simple, non-invasive, and cost-effective technology for sensing muscle activity. However, EMG is also noisy, complex, and high-dimensional. It has nevertheless been widely used in a host of human-machine-interface applications (electrical wheelchairs, virtual computer mice, prosthesis, robotic fingers, etc.) and in particular to measure reaching and grasping motions of the human hand. Here, we developd a more automated pipeline to predict object weight in a reach-and-grasp task from an open dataset relying only on EMG data. In that we shifted the focus from manual feature-engineering to automated feature-extraction by using raw (filtered) EMG signals and thus letting the algorithms select the features. We further compared intrinsic EMG features, derived from several dimensionality-reduction methods, and then ran some classification algorithms on these low-dimensional representations. We found that the Laplacian Eigenmap algorithm generally outperformed other dimensionality-reduction methods. What is more, optimal classification accuracy was achieved using a combination of Laplacian Eigenmaps (simple-minded) and k-Nearest Neighbors (88% for 3-way classification). Our results, using EMG alone, are comparable to others in the literature that used EMG and EEG together. They also demonstrate the usefulness of dimensionality reduction when classifying movement based on EMG signals and more generally the usefulness of EMG for movement classification.

## Introduction

The neuromuscular activations associated with the contraction potentials of the skeletal muscles generate electrical fields. These can be noninvasively recorded to study the activation of muscles, known as surface electromyogram (EMG) [1]. EMG signals are non-stationary in nature and are affected by the structural and functional characteristics of muscles [2]. EMG signals have been widely used and applied in various industrial and clinical settings [3, 4]. Potential applications for signal classification and control of surface EMG include robotic fingers, electric wheelchairs, multifunction prosthesis, virtual keyboard and mouse, navigation in virtual worlds, and many more [3].

Basic human control tasks, such as reaching and grasping, are important for human interfaces for controlling robotic systems [5–7]. Identification of hand movements based on EMG measurements have been largely used in the field of computer and automatic video games, robotic exoskeleton, operative devices and for power prostheses and, thus has been the subject of many studies over the past few years [8–11]. A large number of these studies focus on feature selection for EMG movement classifications and include a dimensionality-reduction step followed by machine-learning-based classification.

As has become apparent, successful classification and pattern recognition of EMG signals requires three main steps: i) data preprocessing, ii) feature extraction iii) classification. During the past several decades, various manual EMG feature-extraction methods have been explored, in the time and/or frequency domains [12]. Finding optimal feature vectors plays an important role in EMG classification [13, 14]. Feature extraction is a method to find intrinsic and meaningful information that is hidden in EMG signal [13]. Appropriate feature extraction tends to result in high classification accuracy [15]. Many studies explored different methods for feature extraction, though the set of feature vectors often carry a number of redundant features [15, 16].

A common method to extract features from signals is dimensionality reduction. So, we briefly present recent approaches for learning low dimensional embeddings from points in high dimensional spaces [17]. Most of them are inspired by linear techniques, such as Principal Component Analysis (PCA) [18] and Multi-Dimensional Scaling (MDS) [19]. Applying PCA in the high dimensional space allows the extraction of principal components that capture more information than their counterparts in the original space. However, such linear techniques have various limitations when applied to EMG. Linear dimensionality reduction techniques are less reliable and more sensitive to the amount of training samples in the input training set. In addition, linear techniques, by nature model linear relations, which is often not the case for EMG. What is more, linear techniques are global in nature, while non-linear dimensionality-reduction techniques can preserve local structures in the original feature space [20].

Non-linear techniques include Locally Linear Embedding (LLE), which was developed later than PCA and MDA [21]. The LLE algorithm computes the basis of such a low-dimensional space. The dimensionality of the embedding, however, has to be given as a parameter, since it cannot always be estimated from the data [22]. Moreover, the output is an embedding of the given data, but not a mapping from the ambient to the embedding space. LLE is not isometric and often fails by mapping distant points close to each other. Another non-linear technique, ISOMAP, is an extension of MDS that uses geodesic instead of Euclidean distances and thus can be applied to non-linear manifolds [23]. The geodesic distances between points are approximated by graph distances. Then, MDS is applied on the geodesic distances to compute an embedding that preserves the property of points to be close or far away from each other. Here we used the Laplacian Eigenmaps algorithm, which was developed by Belkin and Niyogi [24]. It computes the normalized graph Laplacian of the adjacency graph of the input data, which is an approximation of the Laplace-Beltrami operator on the manifold. It exploits locality-preserving properties that were first observed in the field of clustering. The Laplacian Eigenmaps algorithm can be viewed as a generalization of LLE, since the two become identical when the weights of the graph are chosen according to the criteria of the latter. Much like LLE, the dimensionality of the manifold also has to be provided, the computed embeddings are not isometric, and a mapping between the two spaces is not produced. Just a few studies investigated EMG-based hand movement classification using non-linear dimensionality reduction techniques [20]. Such techniques were applied more commonly to EMG during human gait [25, 26], which generally has less degrees of freedom than human reaching and grasping.

In this study, we used the WAY_EEG_GAL open public dataset, which is freely available (see Materials and Methods) and is commonly used to test techniques to decode sensation and action from EMG in humans performing a grasp-and-grasp task. Twelve participants performed a series of lifting movements, and we attempted to predict the weight of the object (165, 330, or 660 g) only from time-domain EMG data. After preprocessing, the procedure typically begins with a feature extraction step, which may be followed by the application of a dimensionality reduction technique. The obtained reduced features are input into a machine-learning classifier.

We also used a non-linear dimensionality-reduction technique—Manifold Learning [22]—to extract meaningful features for classification. However, to the best of our knowledge, automatic feature-extraction directly from the (filtered) raw EMG time-domain signal has not been attempted before. By directly manipulating raw EMG signals, our study therefore shifts the focus from the manual (human-based) feature engineering to completely automated feature-learning [27–29].

A key objective of this study is to compare between the performances of various linear and non-linear dimensionality-reduction techniques on reach-and-grasp EMG data from the arm and hand as the first step before classification. We also wanted to investigate to what degree would automatically extracting the features from raw (filtered) EMG signals directly would lend itself to good dimensionality reduction and classification. Last, previous work on the WAY_EEG_GAL dataset includeed either EEG alone or EEG and EMG. We wanted to see to what extent we could classify the weights in this reach-and-grasp task using EMG alone. Our results would therefore apply to EMG signal classification in engineering, medicine, neuroscience, and so on.

## Materials and Method

Our methodology for EMG signal classification is illustrated in Fig 1 and detailed below. Briefly, EMG signals were first preprocessed and segmented into the first 8 s before feature extraction. This segmentation ensured that the subject started from home position and returned to the home position, removing noise after returning to the home position (Fig 2). The components corresponding to the highest eigenvalues from the output of the dimensionality-reduction algorithms were extracted as the dominant features. Thereafter, these intrinsic features were used for classification.

**Fig 1.**
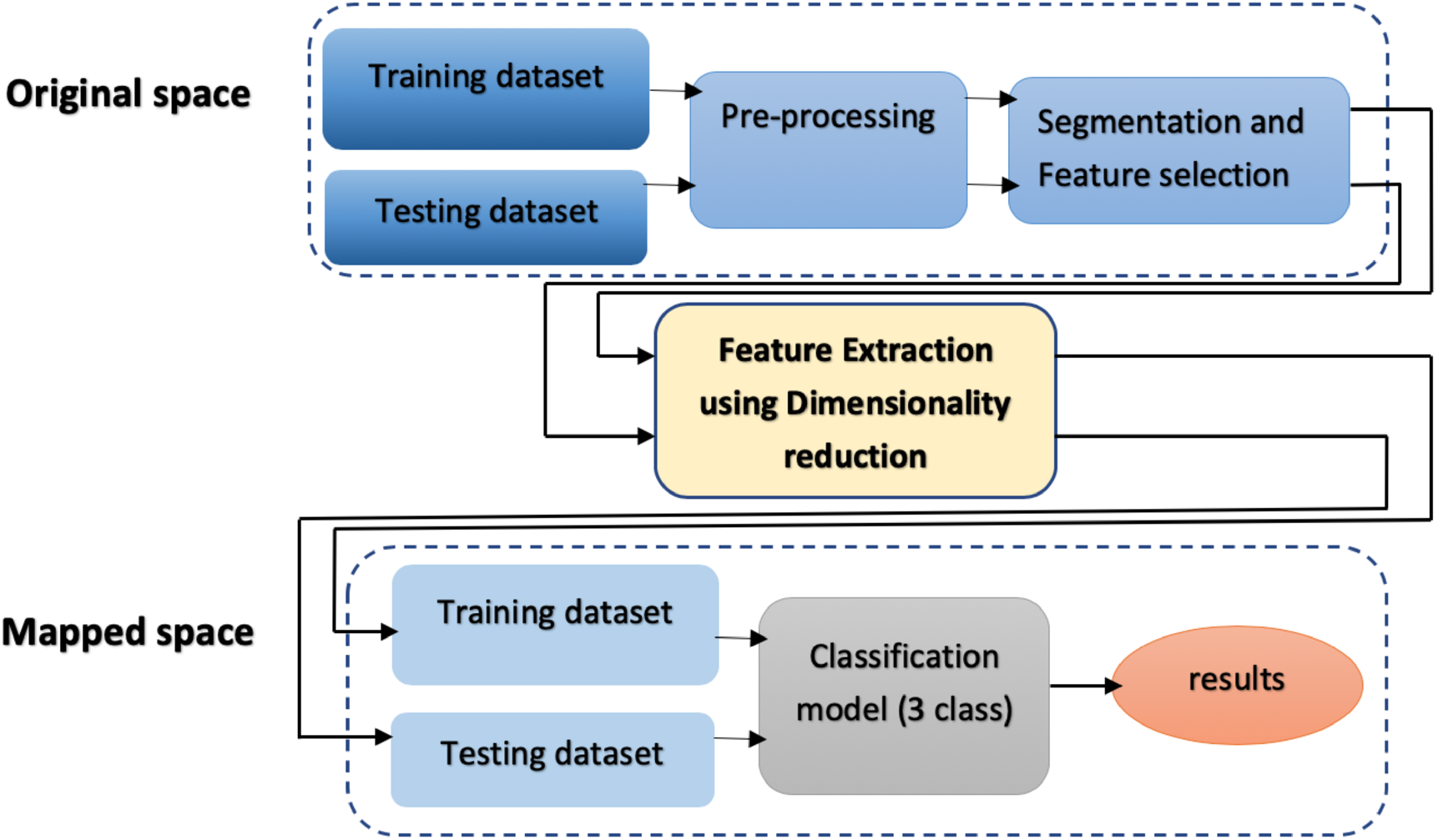
Processing pipeline for EMG signal classification.

**Fig 2.**
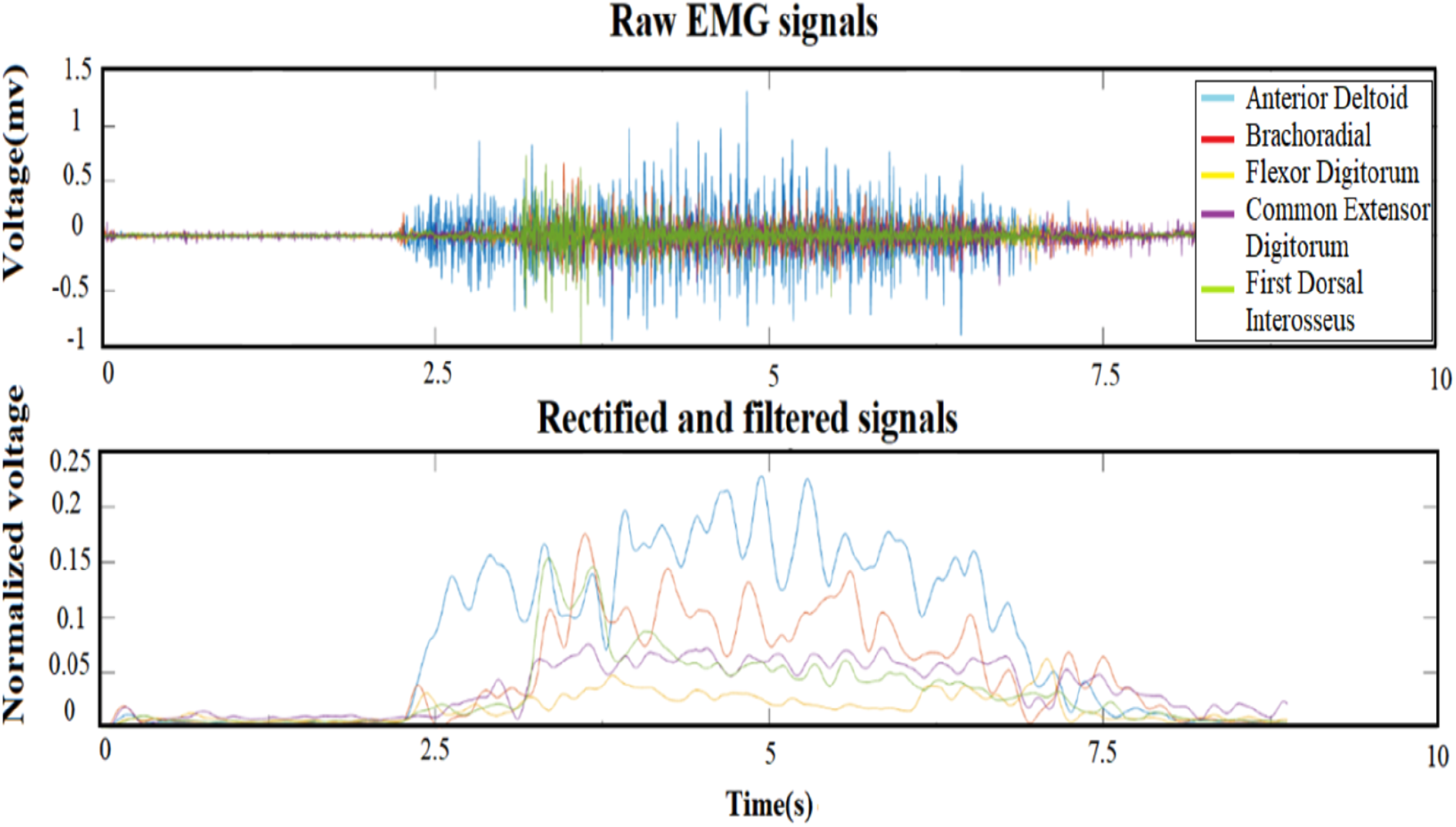
EMG preprocessing. (Top) Raw EMG signals of 5 muscles (Anterior Deltoid, Brachoradial, Flexor Digitorum, Common Extensor Digitorum, and First Dorsal Interosseus). (Bottom) Rectified and filtered EMG signals by band-pass Butterworth filter (4th order) in the 5-450 Hz range on the full-wave rectified and normalized signals from each muscle.

### Dataset

The WAY_EEG_GAL dataset is freely available and has become somewhat of a benchmark to test techniques to decode sensation, intention, and action from surface EMG and scalp EEG in humans performing a reach-and-grasp task (https://doi.org/10.6084/m9.figshare.c.988376) [30]. Here we focus exclusively on the EMG data. Twelve participants performed a series of lifting movements where the object’s weight (165, 330, or 660 g) and surface material (sandpaper, suede, or silk) varied. The EMG signals were sampled at 4 kHz. In each trial, the participants rested their hand in the home position. Then they were cued to reach for the object, grasp it with the thumb and index finger, lift it straight up in the air and hold it for a few of seconds. Then they were instructed to put it back on the support surface, let go of it, and, lastly, return the hand to a designated home position [30]. We used all available 2,645 trials of EMG signals, across all 12 subjects, containing trials with different weights (840 trials for 165 g, 1122 trials for 330 g, and 683 trials for 660 g). The number of trials for each subject is 220 or 221, and the highest imbalance-ratio between classes for each subject is 0.6 (Table 1). The material in all trials was always sandpaper, as per the original design of the experiment [30]. Five EMG electrodes recorded the activity from 5 muscles (Fig 3).

**Table 1.**
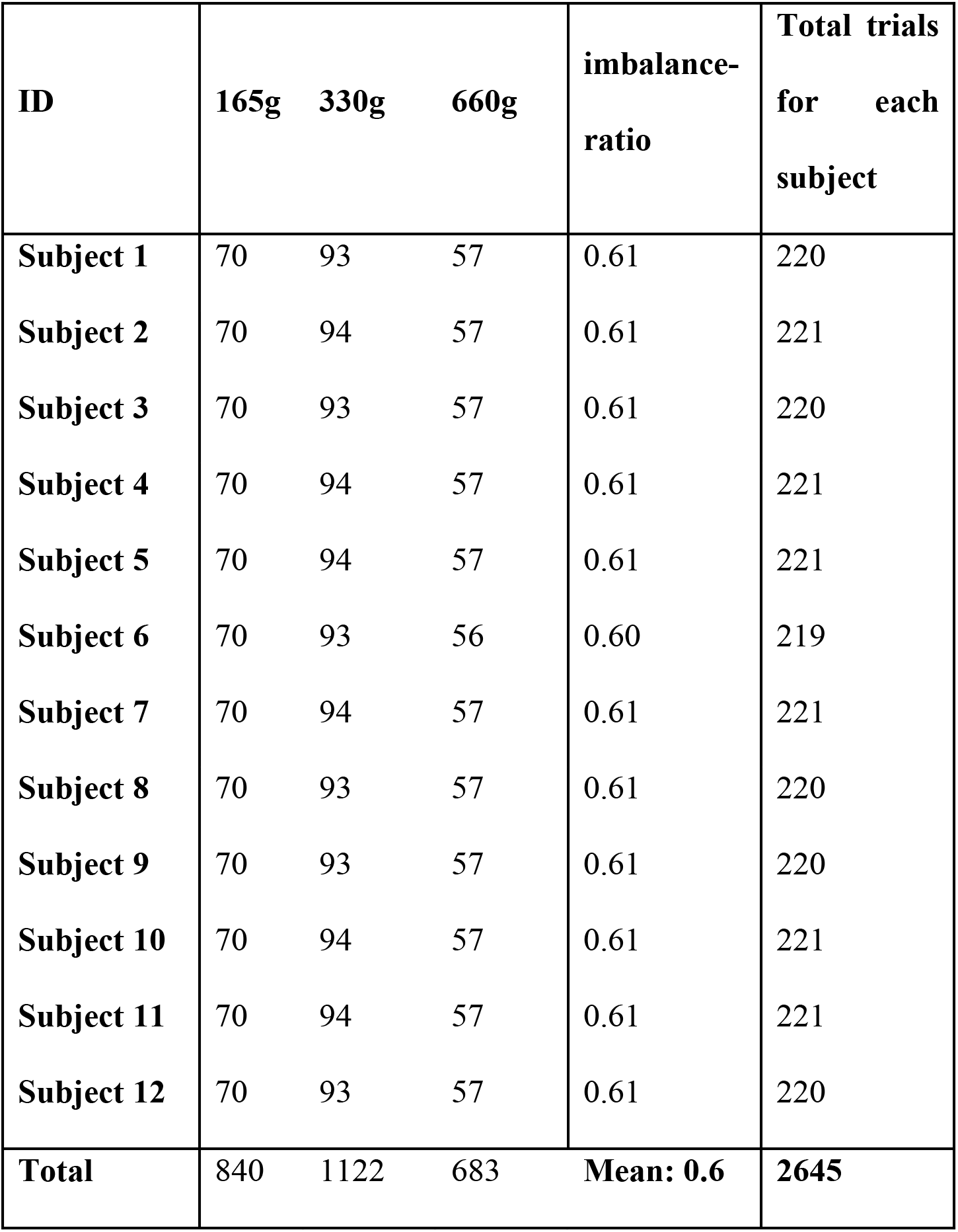
The number of trials for each class and each subject.

| **ID** | **165g** | **330g** | **660g** | **imbalance-**<br>**ratio** | **Total trials**<br>**for each**<br>**subject** |
| --- | --- | --- | --- | --- | --- |
| **Subject 1** | 70 | 93 | 57 | 0.61 | 220 |
| **Subject 2** | 70 | 94 | 57 | 0.61 | 221 |
| **Subject 3** | 70 | 93 | 57 | 0.61 | 220 |
| **Subject 4** | 70 | 94 | 57 | 0.61 | 221 |
| **Subject 5** | 70 | 94 | 57 | 0.61 | 221 |
| **Subject 6** | 70 | 93 | 56 | 0.60 | 219 |
| **Subject 7** | 70 | 94 | 57 | 0.61 | 221 |
| **Subject 8** | 70 | 93 | 57 | 0.61 | 220 |
| **Subject 9** | 70 | 93 | 57 | 0.61 | 220 |
| **Subject 10** | 70 | 94 | 57 | 0.61 | 221 |
| **Subject 11** | 70 | 94 | 57 | 0.61 | 221 |
| **Subject 12** | 70 | 93 | 57 | 0.61 | 220 |
| **Total** | 840 | 1122 | 683 | **Mean: 0.6** | **2645** |

**Fig 3.**
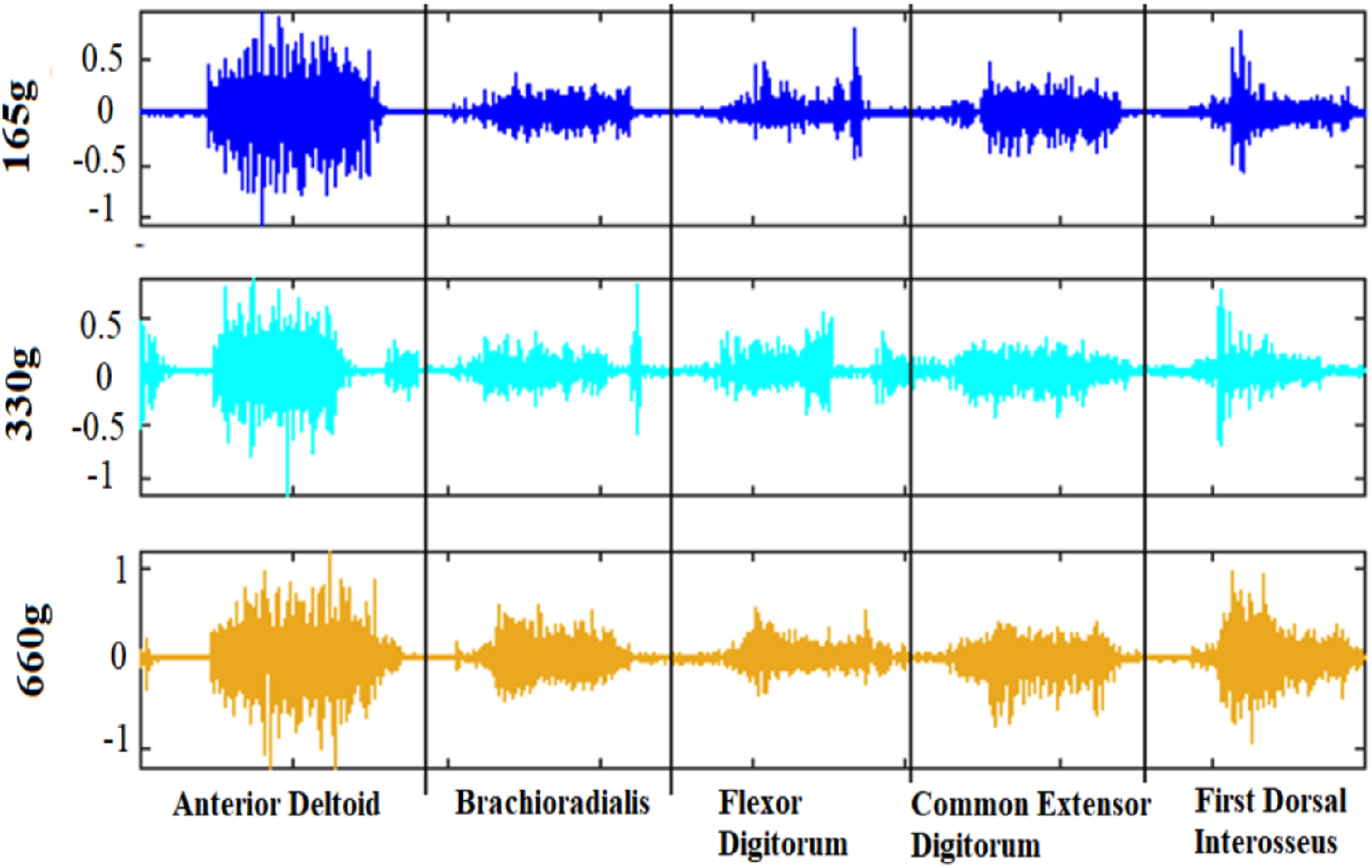
Raw EMG signals of 5 muscles. (Anterior Deltoid, Brachioradialis, Flexor Digitorum, Common Extensor Digitorum, and First Dorsal Interosseous) for 3 different weights (165, 330, and 660 g)

### Preprocessing

All processing below was done on a PC-based (3.4 GHz Intel® CoreTM i7-6700 CPU) using Python 3 and MATLAB 2019b.

The EMG signals are typically contaminated by various types of noise and artifacts. Therefore, preprocessing was necessary and important prior to feature extraction. We used band-pass Butterworth filter (4th order) in the 5-450 Hz range on the full-wave rectified and normalized signals from each muscle (Fig 2).

### Segmentation and feature selection

Since the time required to reach, grasp, and lift is variable for each trial; we consider the first 8 seconds for every trial. This was to remove noise that appeared at the end of the trial, after the subject rested their hand at the home position. For feature selection, we concatenated the signals of the 5 muscles (as in Fig 3). We then subsampled, taking every 5th sample for increased processing speed (lowpass filtering was already carried out as part of the band-pass filter during preprocessing). We ended up with a 5 × 8 × 800 (muscle x time (second) x samples) = 32,000 features.

### Feature Extraction using dimensionality reduction

EMG signals are complex, high-dimensional, and non-linear and hence hard to study in their original form. The effort has therefore been put into finding meaningful, low-dimensional features of these signals. Classical dimensionality-reduction techniques include linear methods such as principal component analysis (PCA) [31] and linear discriminant analysis (LDA) [32]. These techniques preserve the global structure, but at the cost of obscuring local features and preventing any local manipulation of the data. Manifold learning is a technique for recovering a low-dimensional representation from non-linear, high-dimensional data [22, 33]. The literature on manifold learning is dominated by spectral methods. These have a characteristic computational pattern. The first step involves the computation of the k-nearest neighbors (k-NN) of all N data points. Then, an N×N square matrix is populated using some geometric principle. This characterizes the nature of the desired low-dimensional embedding. The eigenvalue decomposition of this matrix is then used to obtain the low-dimensional representation of the manifold.

A trade-off between preserving local and global structures must often be made when inferring the low-dimensional representation. Manifold learning techniques such as Locally Linear Embedding (LLE) [21], Laplacian Eigenmaps [24], t-Distributed Stochastic Neighbor Embedding (t-SNE) [34] are considered to be local methods. This is because they are designed to minimize some form of local distortion and hence result in embedding, which preserves locality. Methods such as ISOMAP [22] are considered global because they enforce the preservation of all geodesic distances in the low-dimensional embedding. All spectral techniques are parameterless (except for neighborhood size; see below) in nature and hence do not characterize the map that generates them. In this study, we compared between different algorithms for manifold learning (Global: ISOMAP; and local: LLE, t-SNE, Laplacian Eigenmaps) and compared them with linear dimensionality-reduction techniques (PCA and LDA). (Note that all the dimensionality-reduction techniques are unsupervised except LDA.)

### The Laplacian Eigenmap algorithm

As we found Laplacian Eigenmaps to be especially successful, hence we describe it in more detail. Given *k* points *x*_1_, …, *x*_*k*_ in *R*^*l*^, find a set of points *y*_1_, …, *y*_*k*_ in ℝ^*m*^ (*m* ≪ *l*) such that *y*_*i*_ represents *x*_*i*_. Therefore, *x*_*i*_, …, *x*_*k*_ ∈ ***M*** and M is a manifold embedded in ℝ^*l*^. The Laplacian Eigenmaps (spectral embedding) is based on the following steps:

**Algorithm 1.**
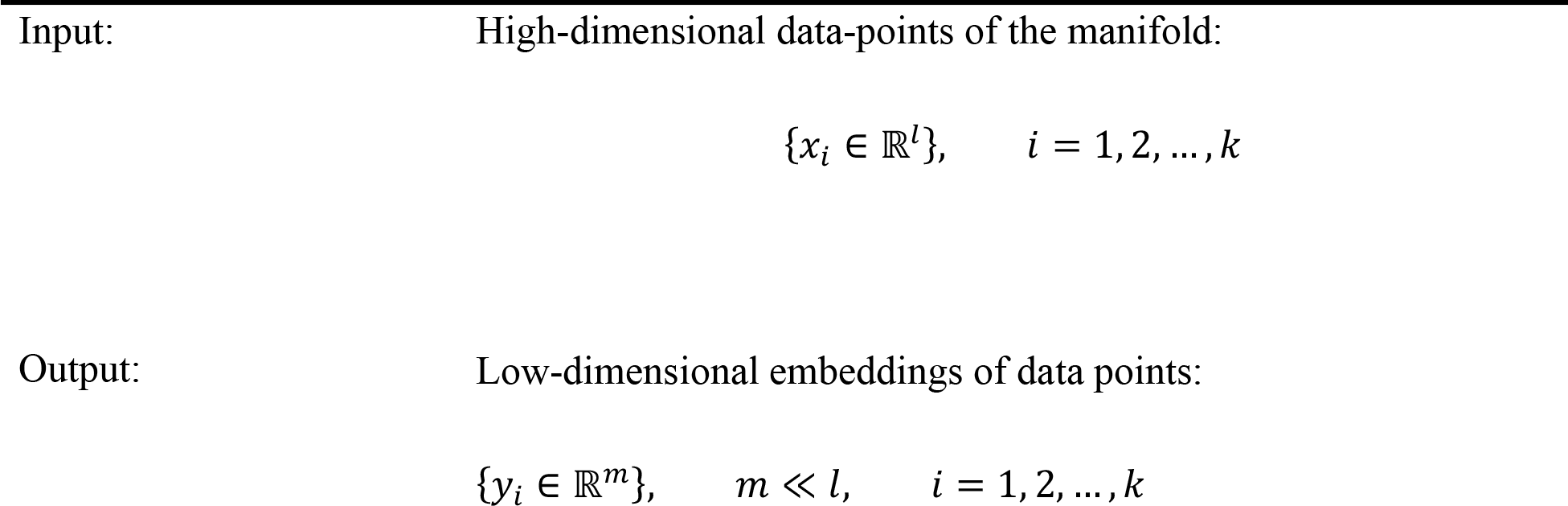
Laplacian Eigenmaps.

Step1. Constructing the graph: We put an edge between nodes *i* and *j* if *x*_*i*_ and *x*_*j*_ are n nearest neighbors. Nodes *i* and *j* are connected by an edge if *i* is among *n* nearest neighbors of *j* or *j* is among *n* nearest neighbors of *i*. We will have connected graph.
Step 2. Choosing the weights. There are two possible way for choosing the weights:

a. *Heat kernel*: 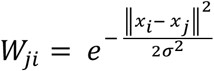 if vertices I and j are connected by an edge and *W*_*ji*_ = 0 if vertices I and j are not connected by an edge (parameter t∈ℝ)
b. *Simple – minded*: *Wij* =

if vertices *i* and *j* are connected by an edge and *W*_*ji*_ =
0 if vertices *i* and *j* are not connected by an edge.
Step 3. Eigenmaps: Compute eigenvalues and eigenvectors for the generalized eigenvector problem:

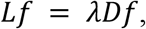 Where D is diagonal weight matrix, and its elements are column (or row, since W is symmetric) sums of W.

*D*_*ii*_ = ∑_*j*_ *W*_*ji*_, *L* = *D* – *W* is the Laplacian matrix Laplacian is a symmetric, positive semidefinite matrix that can be thought of as an operator on functions defined on vertices of *G*.

We leave out the eigenvector corresponding to eigenvalue 0 and use the next *m* eigenvectors for embedding in *m*-dimensional Euclidean space: *x*_*i*_ → *f*_1_(*i*), …, *f*_*m*_(*i*)

The *m* eigenvectors will be considered as features of the dataset.

### Optimal embeddings

Given a data set, we construct a weighted graph G = (V, E) with edges connecting nearby points to each other (assuming the graph is connected). Consider the problem of mapping the weighted graph G to a line so that connected points stay as close together as possible. Let y = (*y*_1_, *y*_2_, …, *y*_*n*_)^*T*^ be such a map. A reasonable criterion for having good mapping is to minimize the following objective function:

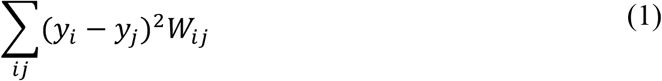

under appropriate constraints. The objective function with our choice of weights *W*_*ij*_ incurs a heavy penalty if neighboring points *x*_*i*_ and *x*_*j*_ are mapped far apart. Therefore, minimizing it is an attempt to ensure that if *x*_*i*_ and *x*_*j*_ are “close,” then *y*_*i*_ and *y*_*j*_ are close as well. It turns out that for any y, we have:

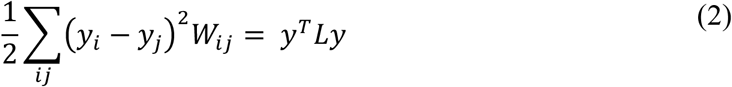

Where as *L* = *D* – *W* and *W*_*ij*_ is symmetric and *D*_*ii*_ = ∑_*j*_ *W*_*ji*_. Thus,

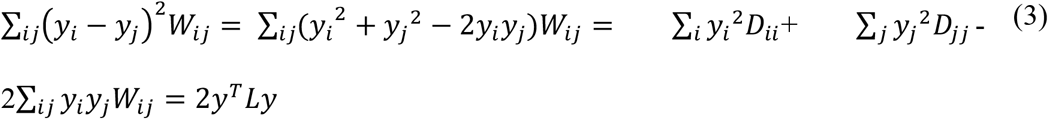

Note that this calculation also shows that L is positive semidefinite. Therefore, the minimization problem reduces to finding: *argmin y*^*T*^ *Ly s. t. y*^*T*^ *Dy* = 1

The constraint *y*^*T*^ *Dy* = 1 removes an arbitrary scaling factor in the embedding. Matrix D provides a natural measure on the vertices of the graph. The bigger the value *D*_*ii*_ (corresponding to the *ith* vertex) is, the more important is that vertex. Because L is positive semidefinite, the vector y that minimizes the objective function is given by the minimum eigenvalue solution to the generalized eigenvalue problem: *Ly* = *λDy*

Let **1** be the constant function taking 1 at each vertex. It is easy to see that **1** is an eigenvector with eigenvalue 0. If the graph is connected, **1** is the only eigenvector for λ = 0. To eliminate this trivial solution, which collapses all vertices of G onto the real number 1, we put an additional constraint of orthogonality and look for: *argmin y*^*T*^ *Ly s. t. y*^*T*^ *Dy* = 1 and *y*^*T*^ *D***1** = 0

Thus, the solution is now given by the eigenvector with the smallest nonzero eigenvalue. The condition *y*^*T*^ *D***1** = 0 can be interpreted as removing a translation invariance in y.

Now consider the more general problem of embedding the graph into m-dimensional Euclidean space. The embedding is given by the *k* × *m* matrix ϒ = [*y*_1_, *y*_2_, …, *y*_*m*_], where the *i*^*th*^ row provides the embedding coordinates of the *ith* vertex. Similarly, we need to minimize:

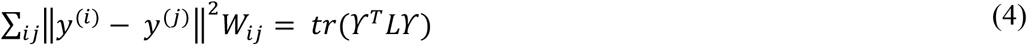

where *y*^(*i*)^ = [*y*_1_(*i*), …, *y*_*m*_(*i*)]^*T*^ is the m-dimensional representation of the i^th^ vertex. This reduces to finding:

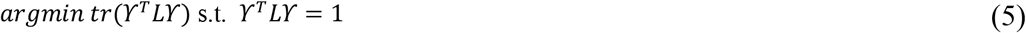

For the one-dimensional embedding problem, the constraint prevents collapse onto a point. For the m-dimensional embedding problem, the constraint presented above prevents collapse onto a subspace of dimension less than *m* – 1.

The above therefore suggest that the Laplacian Eigenmap algorithm keeps samples from the original, higher-dimensional space close to each other also in the lower-dimensional embedding.

### Smooth functions on the manifold

The Laplacian of a graph is analogous to the Laplace Beltrami operator on manifolds. The eigenfunctions of the Laplace Beltrami operator have properties desirable for embedding [24, 35]. Let 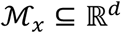 be a smooth, compact manifold embedded in a d-dimensional Euclidean space. A function 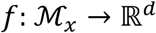 is said to be smooth if 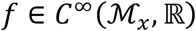, that is, the function *f* and all of its derivatives are continuous. In this section, we define a different notion of smoothness related to the Laplace-Beltrami operator. The Laplace-Beltrami operator *L*_*x*_ is a linear operator generalizing the Laplacian on Euclidean spaces to Riemannian manifolds. The eigenfunctions *ψ*_*i*_ of the Laplace-Beltrami operator span a dense subset of the function space 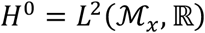. The eigenvalues of the Laplace-Beltrami are real (and non-negative), so we can sort the associated eigenfunctions *ψ*_*i*_ such that *λ*_*i*_ ≤ *λ*_*j*_ for *i* < *j*. We say that *ψ*_*i*_ is smoother than *ψ*_*j*_ if *λ*_*i*_ ≤ *λ*_*j*_. It was shown that the best representation basis, in terms of truncated representation of functions 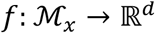 such that ∥ *∇f* ∥ ≤ 1, are in fact the eigenfunctions of the Laplace-Beltrami operator *L*_x_. Thus, in that sense, we say that 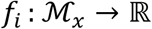 with ∥ *f*_*i*_ ∥ = 1 is smoother than 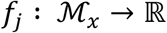 with ∥ *f*_*j*_ ∥ = 1 if ∥*L*_x_*f*_*i*_ ∥≤∥*L*_x_*f*_*j*_ ∥.

Heat kernels and the choice of weight matrix: The Laplace Beltrami operator on differentiable functions on a manifold 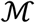 is intimately related to the heat flow. Let *f*: *M* ∶, → ℝ be the initial heat distribution and *u*(*x*, *t*) be the heat distribution at time *t* (*u*(*x*, 0) = *f* (*x*)) (see [24, 35] for more details). The results show that we compute the graph Laplacian with the following weights:

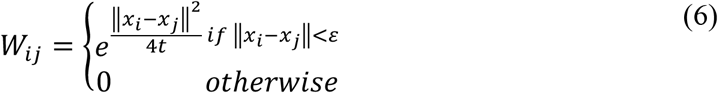

The low dimensional representation of data set that optimally preserves local neighborhood information may be viewed as a discrete approximation to a continuous map that naturally arises from the geometry of the manifold. It is worth highlighting some aspects of Laplacian Eigenmaps here: 1) The algorithm reflects the intrinsic geometric structure of the manifold, which is simple with few local computations and one sparse eigenvalue problem. 2) The justification for the algorithm comes from the role of the Laplace Beltrami operator in providing an optimal embedding for the manifold. The key role of the Laplace Beltrami operator in the heat equation that enables us to use the heat kernel to choose the weight decay function in a in a principled manner. Thus, the embedding maps for the data approximate the Eigenmaps of the Laplace Beltrami operator, which are maps that intrinsically depend on the entire manifold. 3) The locality preserving character of the Laplacian Eigenmap algorithm makes it relatively insensitive to outliers and noise. Close connections to spectral clustering algorithms developed in learning and computer vision. To help gain intuition about manifold-learning algorithms, we demonstrate their use on a simple, spherical dataset (2000 random points on the surface of a 3D sphere) and “Swiss roll” (The 2000 points chosen at random from the Swiss roll; Fig 4). We used the Scikit-learn Python package [36] and Matlab drtoolbox [37, 38] for dimensionality reduction. Laplacian Eigenmaps are termed Spectral Embedding (SE) in Scikit-learn. It has 2 different methods (heat kernel and simple-minded) for constructing the weight matrix. The kernel function for Heat-kernel 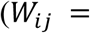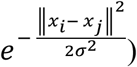 in this package is a Gaussian radial basis function kernel (RBF) with 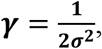, where γ is a parameter that sets the “spread” of the kernel. The results of manifold-learning techniques for 8 neighbors in 2D space are shown in Fig 4. Laplacian Eigenmaps (simple-minded) is the fastest algorithm: the computation time for SE-rbf is 5.45 times longer for a sphere and 31.7 times longer for a Swiss roll. It appears that the construction of the weight matrix drives this difference in computation time.

**Fig 4.**
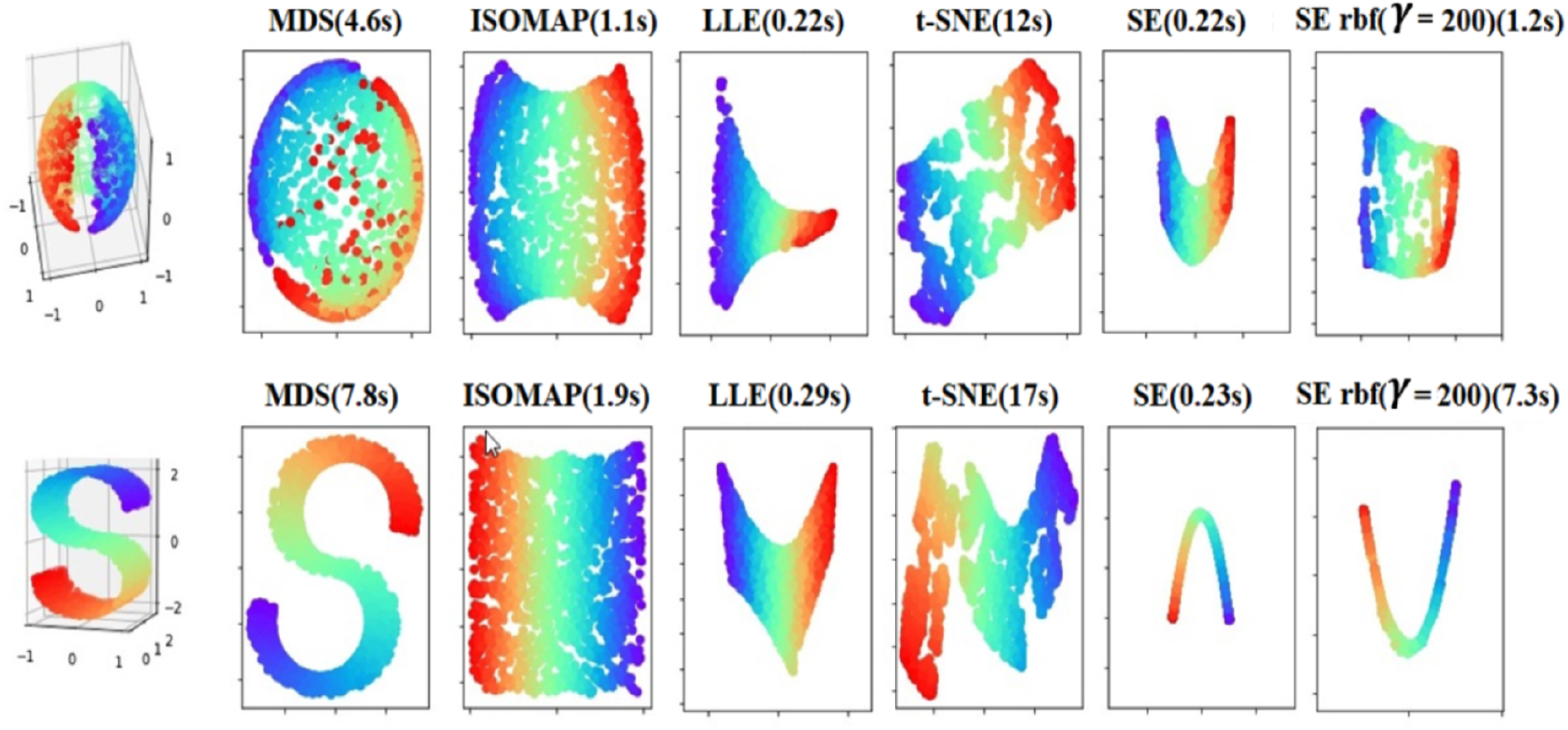
Manifold learning techniques. MDS, ISOMAP, LLE, Spectral embedding (SE) or Laplacian Eigenmaps, and t-SNE on 2000 points randomly distributed on the surface of a sphere. The number of seconds after each method’s name represents its computation time. The first column for SE is simple-minded constructing weight matrix and last column is for heat kernel.

In Table 2, the dimensionality reduction techniques are listed by four general properties: (1) the parametric nature of the mapping between the high-dimensional and the low-dimensional space, (2) the main free parameters that have to be optimized, (3) the computational complexity of the main computational part of the technique, and (4) the memory complexity of the technique [37, 38]. Table 2 shows that most techniques for dimensionality reduction are non-parametric. This means that the technique does not specify a direct mapping from the high-dimensional to the low-dimensional space (or vice versa). The non-parametric nature of most techniques is a disadvantage for two main reasons: (1) it is not possible to generalize the mapping to a held-out or to a new test data (without performing the dimensionality reduction technique again); (2) it is difficult to obtain insights into how much information of the high-dimensional data was retained in the low-dimensional space by reconstructing the original data from the low-dimensional representation of the data and measuring the error between the reconstructed and true data. Regarding free parameters property, Table 2 shows that the objective functions of most non-linear techniques for dimensionality reduction have free parameters that need to be optimized. In other words, there are parameters that directly influence the optimized cost function. Non-convex techniques for dimensionality reduction have additional free parameters, such as the learning rate and the permitted maximum number of iterations. Moreover, LLE uses a regularization parameter in the computation of the reconstruction weights. The presence of free parameters has both advantages and disadvantages. The main advantage of the presence of free parameters is that they make the technique more flexible. However, they then need to be tuned to optimize the performance of the dimensionality reduction technique. Table 2 also provides more details in computational and memory complexities of the techniques. The computational complexity of a dimensionality-reduction technique is important for its practical applicability. Algorithms grow increasingly infeasible as computational demands or memory rise. The computational complexity of a dimensionality reduction technique is determined by: (1) properties of the dataset, such as the number of datapoints n and their dimensionality D, and (2) by parameters of the techniques, such as the target dimensionality d, the number of nearest neighbors k, σ (for techniques based on neighborhood graphs). In Table 2, p denotes the ratio of nonzero elements in a sparse matrix to the total number of elements.

**Table 2.**
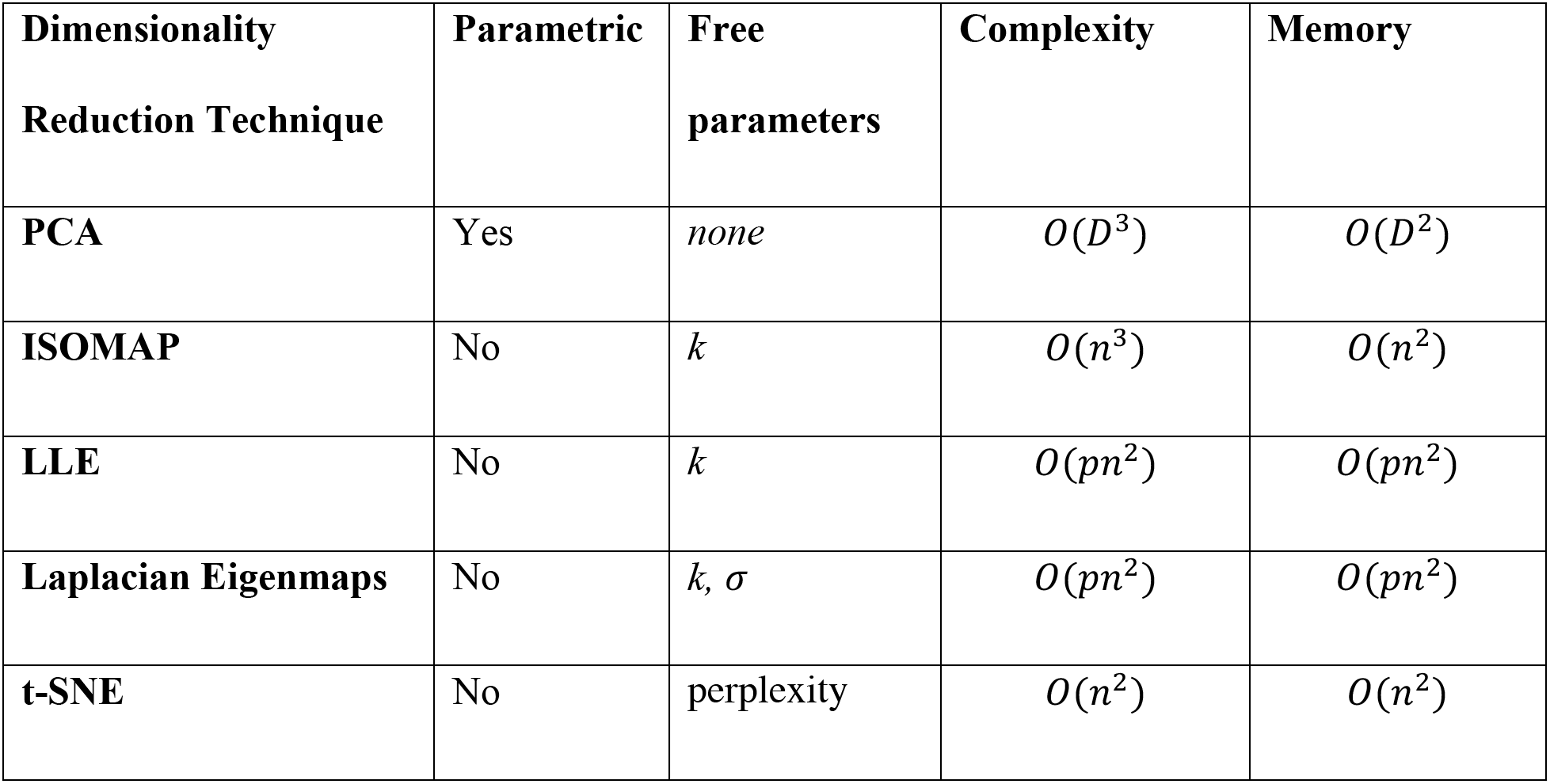
Properties of techniques for dimensionality reduction.

| **Dimensionality**<br>**Reduction Technique** | **Parametric** | **Free**<br>**parameters** | **Complexity** | **Memory** |
| --- | --- | --- | --- | --- |
| **PCA** | Yes | *none* | $O(D^3)$ | $O(D^2)$ |
| **ISOMAP** | No | $k$ | $O(n^3)$ | $O(n^2)$ |
| **LLE** | No | $k$ | $O(pn^2)$ | $O(pn^2)$ |
| **Laplacian Eigenmaps** | No | $k, \sigma$ | $O(pn^2)$ | $O(pn^2)$ |
| **t-SNE** | No | perplexity | $O(n^2)$ | $O(n^2)$ |

### Out-of-sample extension

An important requirement for dimensionality reduction techniques is the ability to embed new high-dimensional datapoints into an existing low-dimensional data representation. However, there is no an explicit projection function between the original data and their low dimensional representations in the original LE algorithm, which makes out-of-sample extension difficult. To find projection of any additional samples, LE needs to be run on all the data together with the additional samples, resulting in considerable computational cost especially when applying it to large scale data pattern recognition. Fortunately, various methods have been developed to mitigate the out-of-sample problem [39]: Linear approximation to LE, Kernel extensions to LE, Tensor representation of LE, incremental learning for LE, neural network approaches, and Extreme Learning Machine. The out-of-sample extension for spectral techniques has been presented in [40]. Further, spectral techniques such as ISOMAP, LLE, and Laplacian Eigenmaps support out-of-sample extensions via the Nyström approximation.

### Nyström extension

Let D denote the dimension of the initial set, N the number of samples (or points), *x*_*i*_ a sample in ℝ^*D*^ and *X* the *D* × *N* training matrix containing the samples. Let *y*_*i*_ denote the coordinates in the embedded space, included in ℝ^*d*^ where B is the reduced dimension that corresponds to *x*_*i*_. Finally, let *x*_*N*+1_ denote a sample not belonging to the initial set of samples, i.e. an out-of-sample point; the goal is to estimate its reduced coordinates *y*_*N*+1_. The Nyström method speeds up kernel-method computations by performing the eigen-decomposition on a subset of examples [41]. It was previously used to propose an out-of-sample extension to kernel-based spectral methods [40]. Let us recall the general framework in which spectral dimension reduction techniques can be cast. Let *W* be a symmetric matrix of size *N* × *N*, expressing the affinity between the N points of the training set. Let *K*(·,·) denote a data-dependant kernel function giving rise to matrix *W* with *W*_*ij*_ = *K*(*x*_*i*_, *x*_*j*_).

Let (*v*_*k*_, *λ*_*k*_) denote the eigenvector and eigenvalue pairs such that *Wv*_*k*_ = *λ*_*k*_ *v*_*k*_. For dimensionality reduction, retain the *d* largest (or smallest, depending on the method) eigenvalues and their associated eigenvectors. The embedding (or reduced coordinates) of each training sample *x*_*i*_ is the ith row of a matrix *U* that contains the d eigenvectors in columns. The Nyström extension for an out-of-sample point is a weighted sum of the previously calculated eigenvectors and eigenvalues. More precisely, the kth reduced coordinate of the out-of-sample point is approximated as: 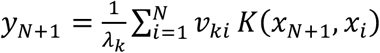 for all *k* = 1, …, B or, in matrix form: 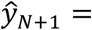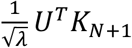, where 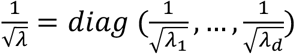. *U* is the matrix whose columns are the eigenvectors, and *K*_*N*+1_ = [*K*(*x*_*N*+1_, *x*_1_) … *K*(*x*_*N*+1_,*X*_*N*_)]. In [4], Bengio et al. have designed a formulation of *K* (·,·) for Laplacian eigenmaps:

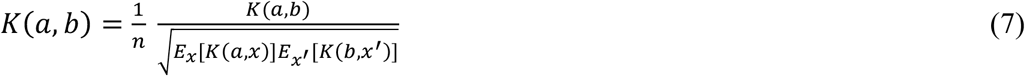

The Nyström extension is applicable to any technique that make use of a kernel function. This method requires some parameter choice for the kernel *K* (·,·), usually made heuristically. In Fig 5 we depict the embedding of additional, out-of-sample points (there termed “test dataset”). As is apparent, the out-of-sample points are mapped to plausible locations in the low-dimensional space.

**Fig 5.**
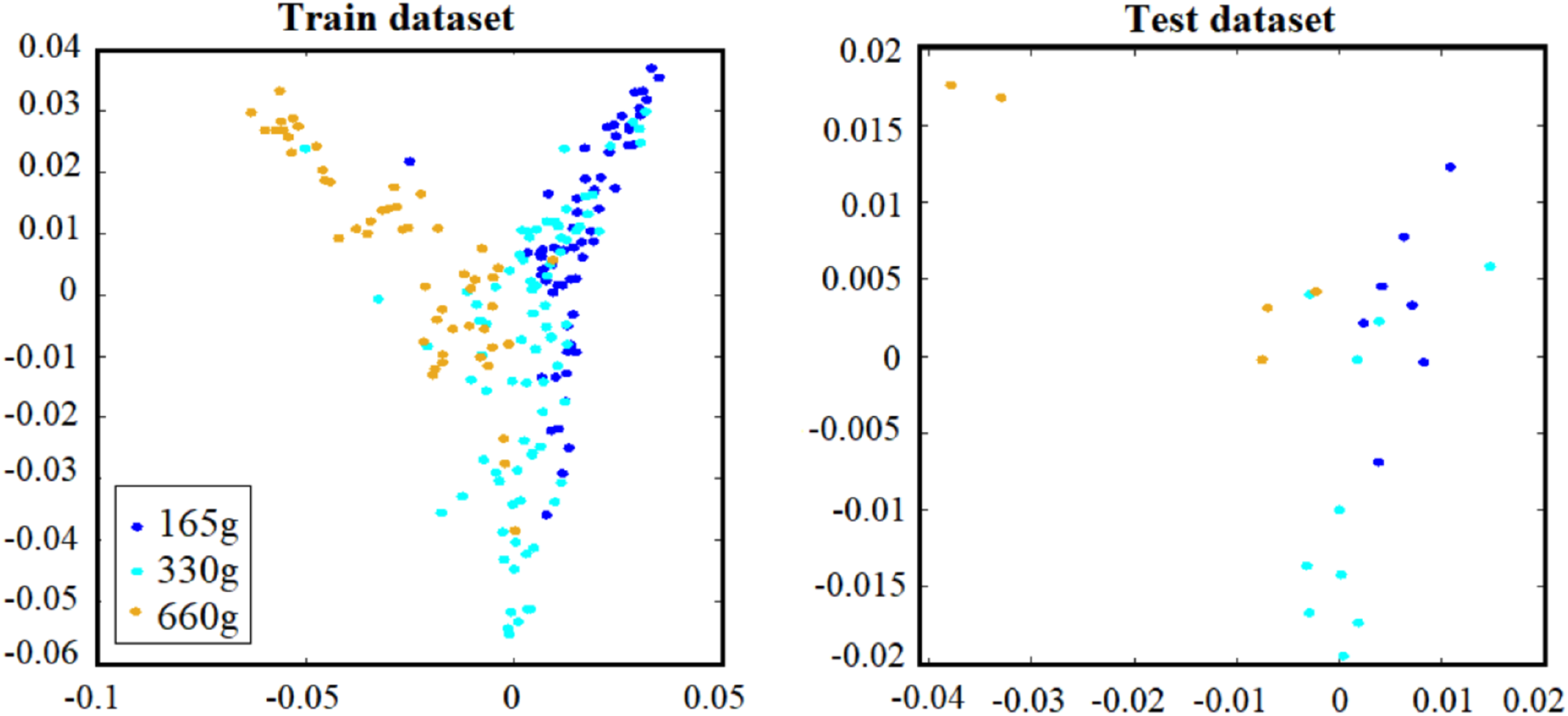
Example of an out-of-sample extension via the Nyström approximation in embedded space by Laplacian Eigenmaps with k = 8 and σ = 1. The training */* test percentage was 90% / 10%.

### Classification

After finding an optimal feature set, we tested commonly used classification algorithms: k-NN [42], linear and RBF SVM [28] (C=32, *γ* = 0.01 for RBF SVM), and Random Forest [43]. We evaluated the performance of each classifier on data after running the above dimensionality-reduction techniques. The dataset was divided into disjoint training and testing sets, which consisted of 90% and 10% of the total trials. Evaluations on the training set relied on 10-fold cross-validation.

Another common method for EMG classification is deep learning [44]. We tested several deep learning architectures on our selected feature set. However, we ran into severe overfitting issues, resulting in accuracies very close to chance level. This is likely due to the relatively small features-to-samples ratio for our dataset. We, therefore, did not include deep-learning results in our analyses.

As the dataset was imbalanced (the highest imbalanced ratio is 0.6), we used the F1-score as a metric of accuracy [45]. The F1-score provides a way to combine both precision and recall into a single measure that captures both properties. Once precision and recall have been calculated for a binary or multiclass classification problem, the two scores can be combined into the calculation of the F1-score.

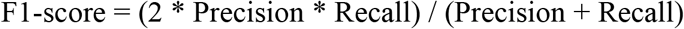

This is the harmonic mean of the two fractions. The F1-Score might be the most common metric used on imbalanced classification problems [46].

## Results

### Parameter settings

The proposed framework has 2 parameters: the number of nearest neighbors in Laplacian Eigenmaps to construct the Laplacian matrix (either using the direct number of neighbors, k, or using a heat kernel approach, **σ**); the number of eigenvectors used for data mapping, that is the dimensions of the mapped space. The number of nearest neighbors in Laplacian eigenmaps, k, is tuned from {4, 5, …, 20}. We checked values of **σ** from {0.1, 1, 10, 100,1000}. Fig 6 shows the effect of different k and **σ** on the training dataset for Subject1. The number of eigenvectors is tuned from **{**1,5, …, length (training trial)}.

**Fig 6.**
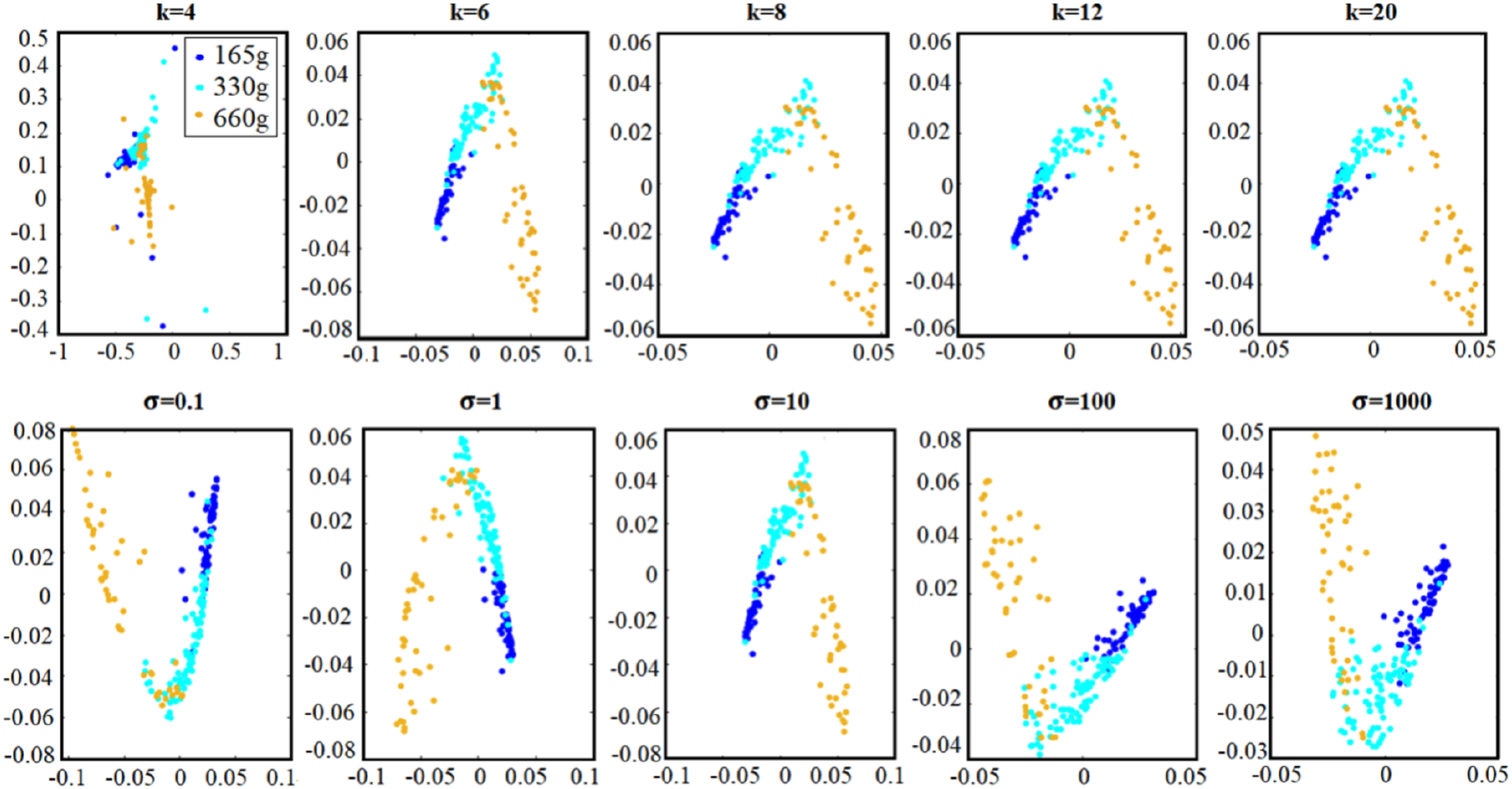
Visualization of the effect of k (Top row) and **σ** (Buttom row) on the training dataset subject1 in the embedded space.

Table 3 demonstrates the best number of eigenvectors for each dimensionality-reduction method for each subject. As a sanity check, we also used the maximum likelihood estimator (MLE) as the intrinsic dimensionality estimator in the Matlab toolbox for dimensionality reduction [38]. The number varies between 82 and 102 over the 12 subjects. Before presenting the quantitative evaluation of classification, it is useful to visualize the embedded EMG data. To this end, we visualized some embedded samples using six methods (PCA, LDA, ISOMAP, LLE, Laplacian Eigenmaps, and t-SNE). Fig 7 shows the 2 prominent components for each of these methods.

**Table 3.**
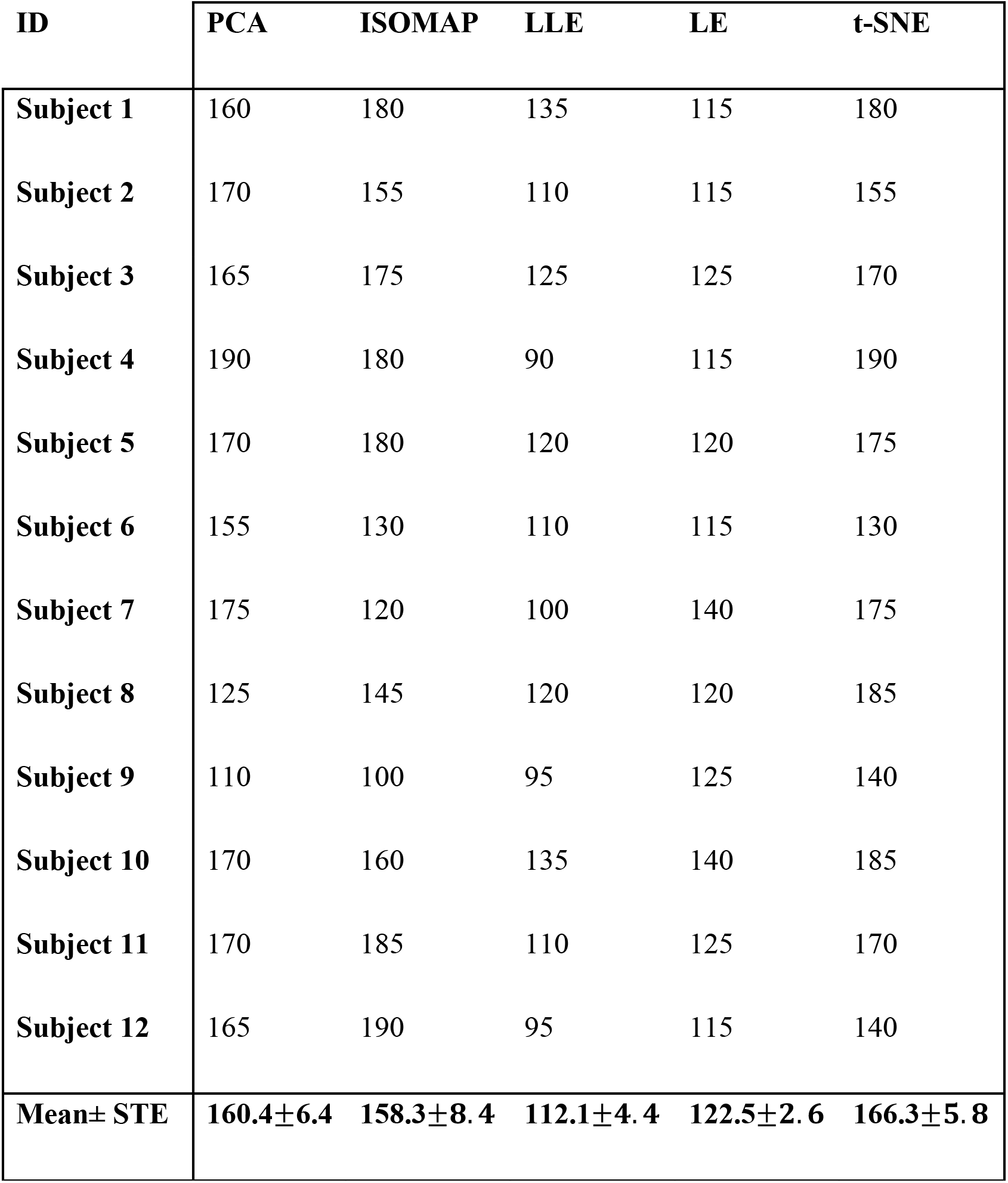
The number of dimensions that leads to the highest F1-score (for k-NN—see Fig 8) vs. different dimensionality reduction techniques for each subject. The mean and standard error (STE) over all subjects for the different methods are also shown.

| **ID** | **PCA** | **ISOMAP** | **LLE** | **LE** | **t-SNE** |
| --- | --- | --- | --- | --- | --- |
| **Subject 1** | 160 | 180 | 135 | 115 | 180 |
| **Subject 2** | 170 | 155 | 110 | 115 | 155 |
| **Subject 3** | 165 | 175 | 125 | 125 | 170 |
| **Subject 4** | 190 | 180 | 90 | 115 | 190 |
| **Subject 5** | 170 | 180 | 120 | 120 | 175 |
| **Subject 6** | 155 | 130 | 110 | 115 | 130 |
| **Subject 7** | 175 | 120 | 100 | 140 | 175 |
| **Subject 8** | 125 | 145 | 120 | 120 | 185 |
| **Subject 9** | 110 | 100 | 95 | 125 | 140 |
| **Subject 10** | 170 | 160 | 135 | 140 | 185 |
| **Subject 11** | 170 | 185 | 110 | 125 | 170 |
| **Subject 12** | 165 | 190 | 95 | 115 | 140 |
| **Mean± STE** | **160.4±6.4** | **158.3±8.4** | **112.1±4.4** | **122.5±2.6** | **166.3±5.8** |

**Fig 7.**
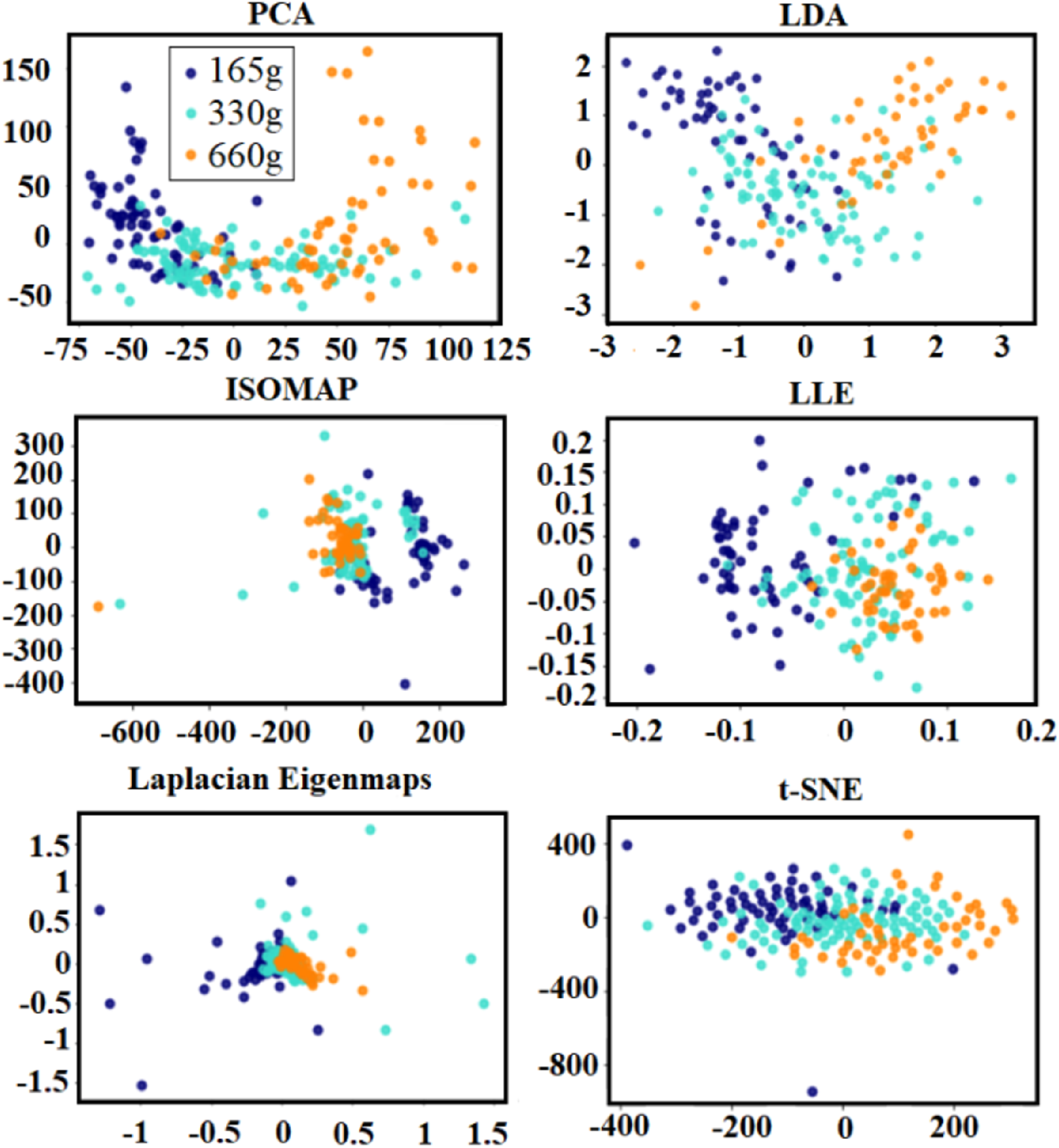
Visualization of the embedding process: EMG visualization using the 2 most prominent components of different dimensionality-reduction technique (The x-axis is the most prominent component and y-axis is the second most prominent component)

### Classifier Performance

We computed the average accuracy using PCA, ISOMAP, LLE, Laplacian Eigenmaps, and t-SNE for the best classifier (k-NN; Fig 8). Since the maximum dimension for LDA is equal to the number of classes minus one, the corresponding curve was not plotted.

**Fig 8.**
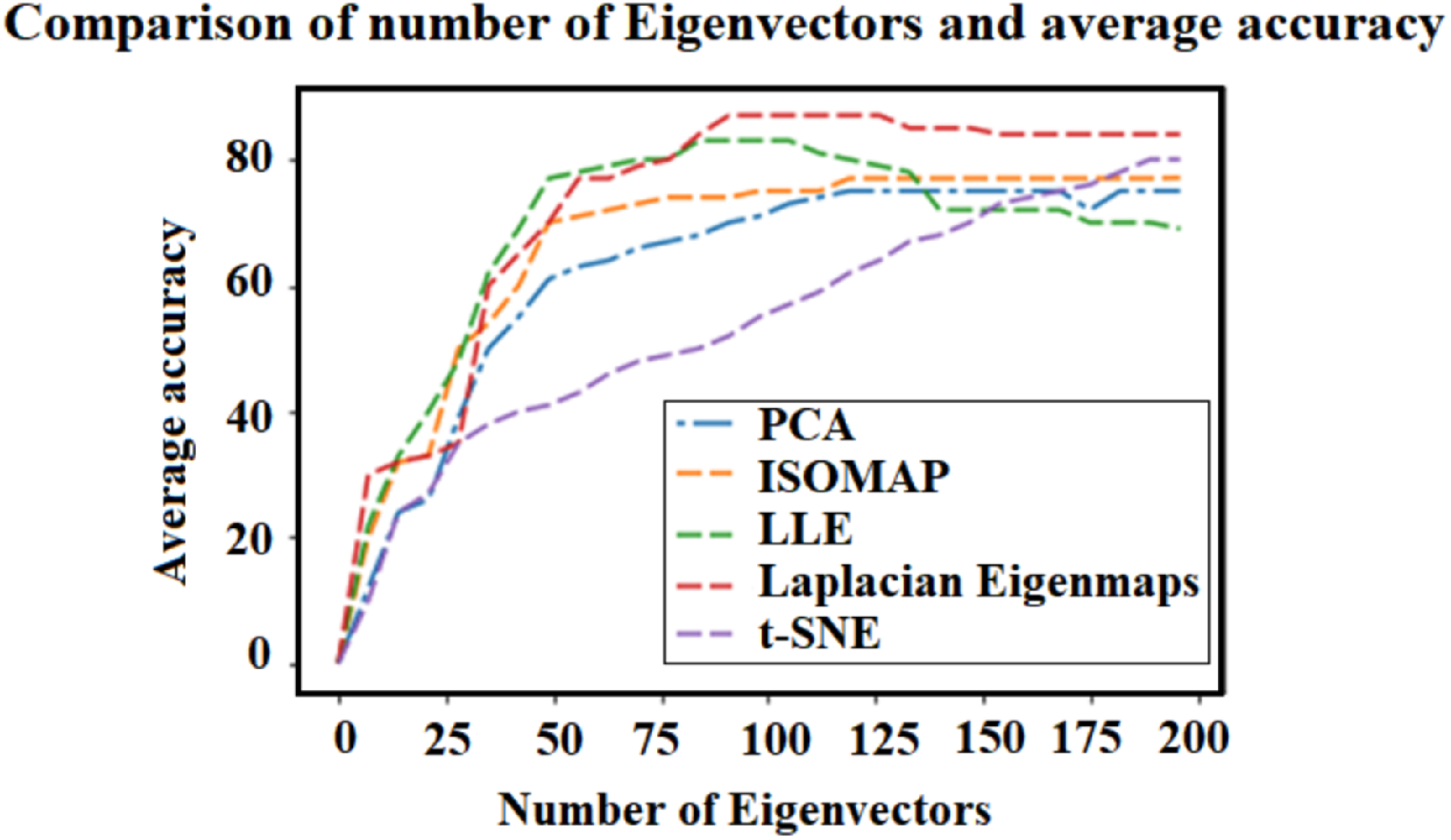
Average accuracy (across all 12 subject) as a function of the number of eigenvectors obtained with different dimensionality reduction techniques for the best classifier (k-NN; see also Table 3).

Table 4 details the prediction accuracies of the different classification algorithms on the test set for the various dimensionality-reduction techniques over all 12 subjects. On average, Laplacian Eigenmaps (especially with heat kernel) is the algorithm with the highest accuracy across all dimensionality-reduction methods—78.15%. And k-NN is the dimensionality-reduction method resulting in the highest mean accuracy across all classification algorithms—80%. Interestingly, their intersection—Laplacian Eigenmaps and k-NN—have the highest overall accuracy, at 88%. In Table 5,the performance of Laplacian Eigenmaps (simple-minded) and different classifiers vs. different number of neighbors (k) is shown. It demonstrates that k=8 fits well for this dataset.

**Table 5.**
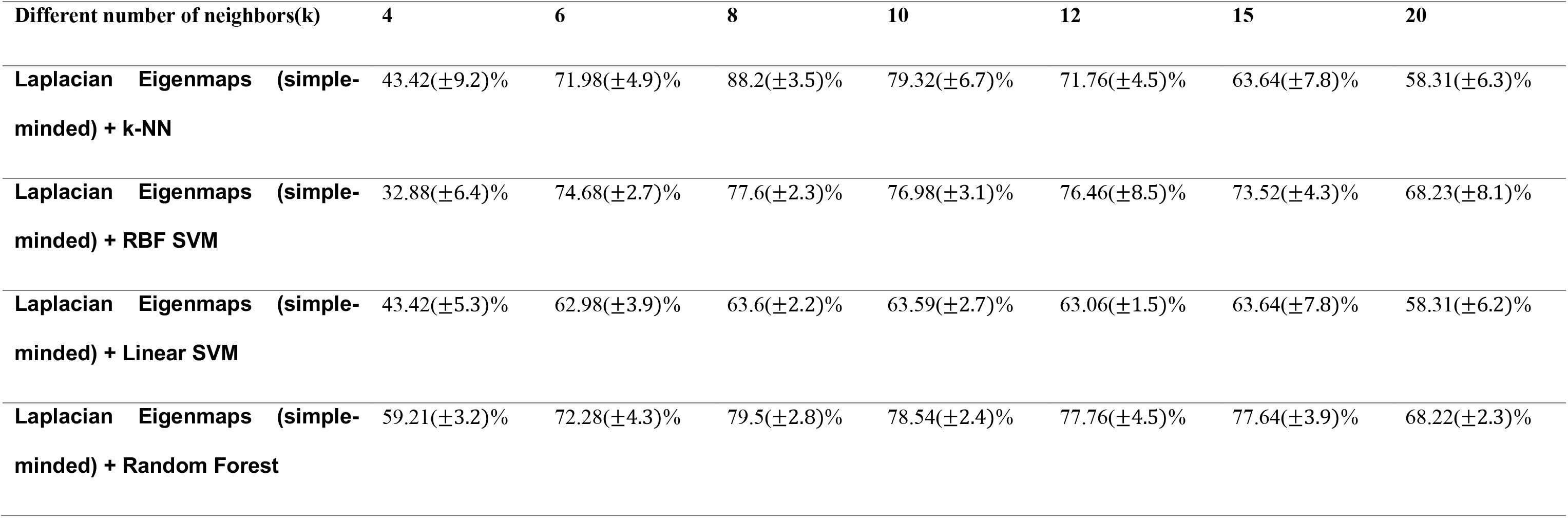
Performance of Laplacian Eigenmaps (simple-minded) and different classifiers vs. different number of neighbors (K) on the EMG signals (accuracy percentage *standard error*)

## Discussion

Our goal in this study was to compare the performances of various linear and non-linear dimensionality-reduction techniques on reach-and-grasp EMG data from the arm and hand as the first step before classification. We worked on raw (filtered) EMG signals directly, automatically extracting the features for the later classification phase. The dimensionality-reduction algorithms we used lowered the dimensionality of our data from 32,000 features to less than 200—so, more than 160-fold. We then applied various classification techniques on this 3-way classification problem and discovered that the combination of Laplacian Eigenmaps (simple-minded, k=8) with the k-NN classifier resulted in the highest classification accuracy (F1 score 88.2±3.5%). To the best of our knowledge, ours is the first successful attempt to use automatic feature-extraction directly from the (filtered) raw EMG time-domain signal [20, 47–52]. What is more, our approach resulted in relatively high decoding accuracy.

Other studies that used the same dataset that we used mostly focused on EEG [47–50]. However, Cisotto et al. used both EEG and EMG to classify the same dataset [51]. They only used 2 of the 3 available classes (the most extreme weights: 165 and 660 gr). They also report their results in terms of accuracy, even though their classes are imbalanced (ratio of 0.81 between the number of trials in the 2 classes). They report a maximal accuracy of 94% (using only the Brachoradial muscle). Running our analysis as is (using all muscles), but with only the 2 weight classes they used, and reporting accuracy instead of F1 score, we get an accuracy of 90.9 ± 2.5%. This accuracy is statistically indistinguishable from theirs (t-test: t(11)=-1.33, p=0.21). Therefore, even though we used only EMG, and we did not focus our analysis on a binary classification problem, we were able to achieve comparable results. This might be due to the superiority of our method— perhaps our automatic feature extraction or our dimensionality-reduction algorithm. Another, not mutually exclusive, possibility is that the high classification accuracy they and we were able to achieve owes more to the EMG signals rather than the EEG signals.

The range of mean accuracies among the dimensionality-reduction algorithms was 70-78% (Table 4). Interestingly, Laplacian Eigenmaps not only performed best on average; its simple-minded version also generally requires the shortest computing time (see Materials and Method). The spread of accuracies across classification algorithms was wider—65 to 80%. It appears that the linearity of linear-SVM was detrimental for EMG signal decoding, while arguably the most non-linear technique, k-NN, faired best.

**Table 4.**
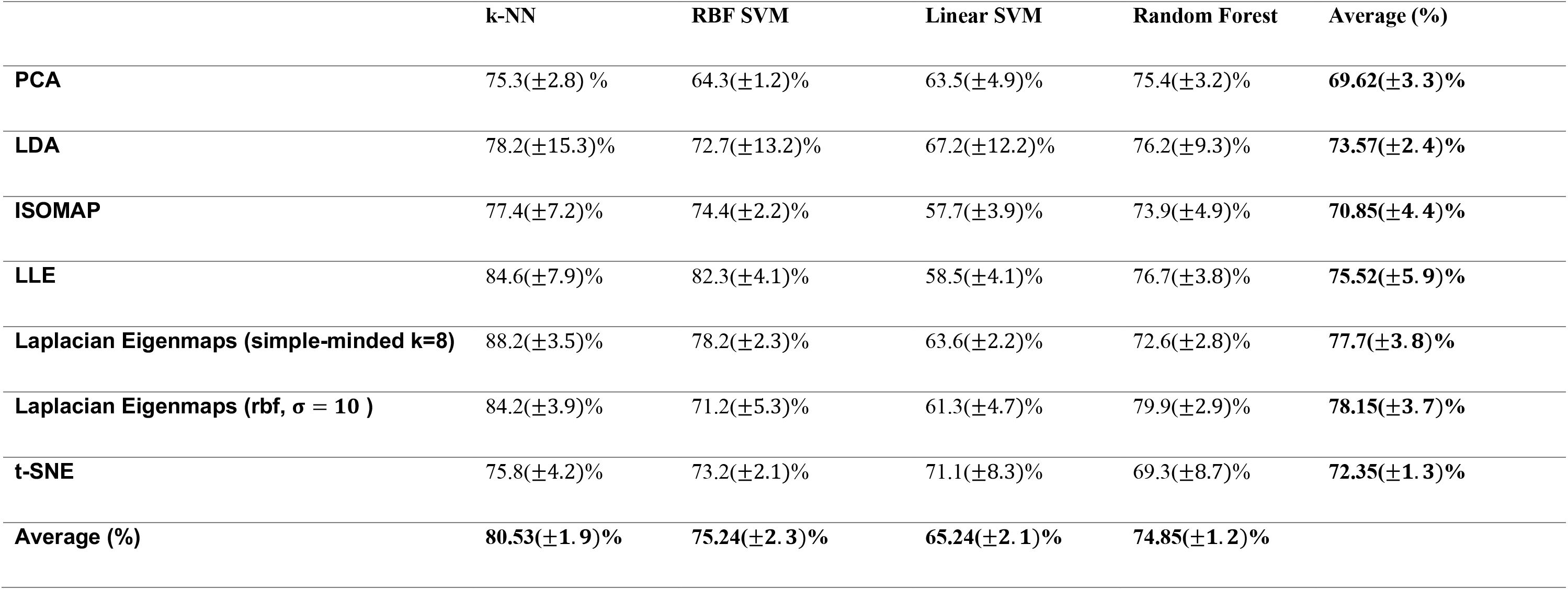
Performance of different classifiers vs. different methods of dimension reduction on the EMG signals (accuracy percentage (±*standard error*))

|  | **k-NN** | **RBF SVM** | **Linear SVM** | **Random Forest** | **Average (%)** |
| --- | --- | --- | --- | --- | --- |
| **PCA** | 75.3(±2.8) % | 64.3(±1.2)% | 63.5(±4.9)% | 75.4(±3.2)% | **69.62(±3.3)%** |
| **LDA** | 78.2(±15.3)% | 72.7(±13.2)% | 67.2(±12.2)% | 76.2(±9.3)% | **73.57(±2.4)%** |
| **ISOMAP** | 77.4(±7.2)% | 74.4(±2.2)% | 57.7(±3.9)% | 73.9(±4.9)% | **70.85(±4.4)%** |
| **LLE** | 84.6(±7.9)% | 82.3(±4.1)% | 58.5(±4.1)% | 76.7(±3.8)% | **75.52(±5.9)%** |
| **Laplacian Eigenmaps (simple-minded k=8)** | 88.2(±3.5)% | 78.2(±2.3)% | 63.6(±2.2)% | 72.6(±2.8)% | **77.7(±3.8)%** |
| **Laplacian Eigenmaps (rbf, $\sigma = 10$ )** | 84.2(±3.9)% | 71.2(±5.3)% | 61.3(±4.7)% | 79.9(±2.9)% | **78.15(±3.7)%** |
| **t-SNE** | 75.8(±4.2)% | 73.2(±2.1)% | 71.1(±8.3)% | 69.3(±8.7)% | **72.35(±1.3)%** |
| **Average (%)** | **80.53(±1.9)%** | **75.24(±2.3)%** | **65.24(±2.1)%** | **74.85(±1.2)%** |  |

It also appears that dimensionality-reduction techniques relying on local embedding were better than those that used global embedding. Such local methods strive to map nearby samples on the original manifold to nearby samples in the low-dimensional space (and vice versa for far away samples). Global methods, in contrast, strive for a faithful representation of the data’s global structure. As reach-to-grasp motion is composed of different phases of movement, local methods may better preserve the varying geometry across phases. Local methods are also computationally more efficient, involving only sparse matrix computations. It may further not be surprising that k-NN works best with local dimensionality-reduction methods. These methods keep nearby samples close to each other, facilitating nearest-neighbor approaches like k-NN.

Our method, therefore, resulted in relatively high accuracy on 3-way classification while maintaining automatic feature extraction. What is more, the methodology proposed in this paper is well suited to real-time operation, potentially in combination with EEG [52], because the computational load in training and testing the model is relatively low.

## Conclusion

This study proposes a complete, automated pipeline for the preprocessing, feature selection, feature extraction, and classification of grasping objects of 3 different weights, with the only input being raw (filtered) EMG data from 5 muscles. Besides showcasing relatively high accuracy (F1 score 88.2±3.5%), our study highlights the importance of properly combining feature selection and classification algorithms.

## Acknowledgements

This publication was made possible in part through the support of a joint grant from the John Templeton Foundation and the Fetzer Institute, and Boston Scientific Corporation. The opinions expressed in this publication are those of the author(s) and do not necessarily reflect the views of the John Templeton Foundation or the Fetzer Institute. This publication was also made possible in part by the support of Boston Scientific Investigator-Sponsored Research Program.

